## Supplementary Material for "Growth and rapid succession of methanotrophs effectively limit methane release during lake overturn"

### Supplementary Figures

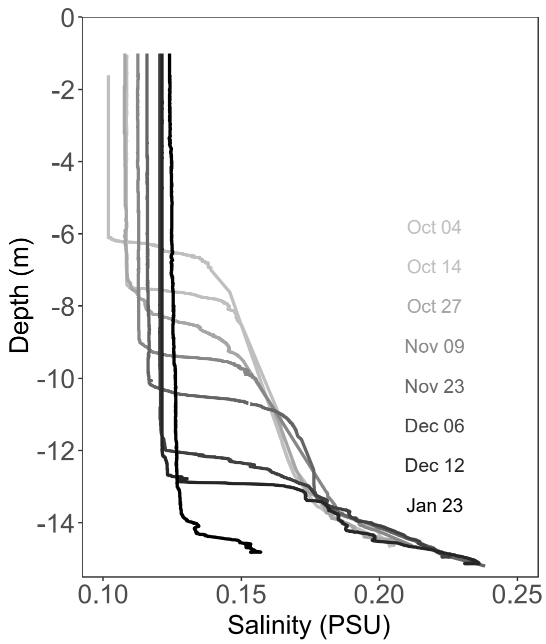


**Supplementary Figure 1** Salinity profiles taken with the profiling in-situ analyser (PIA). Salinity profiles at each sampling date during the lake overturn which continued from October to December. The higher salinity in the bottom water allowed for an inverse stratification on Dec 12 (see Fig. 1B)

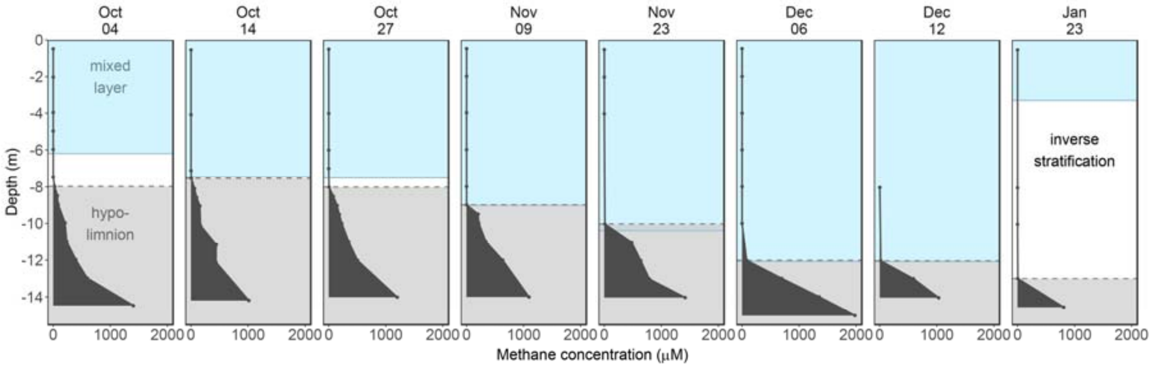


**Supplementary Figure 2** Methane concentration profiles measured at each sampling date. The methane-enriched hypolimnion (≥ 2 µM) is shaded in gray, the mixed layer is shaded in blue.

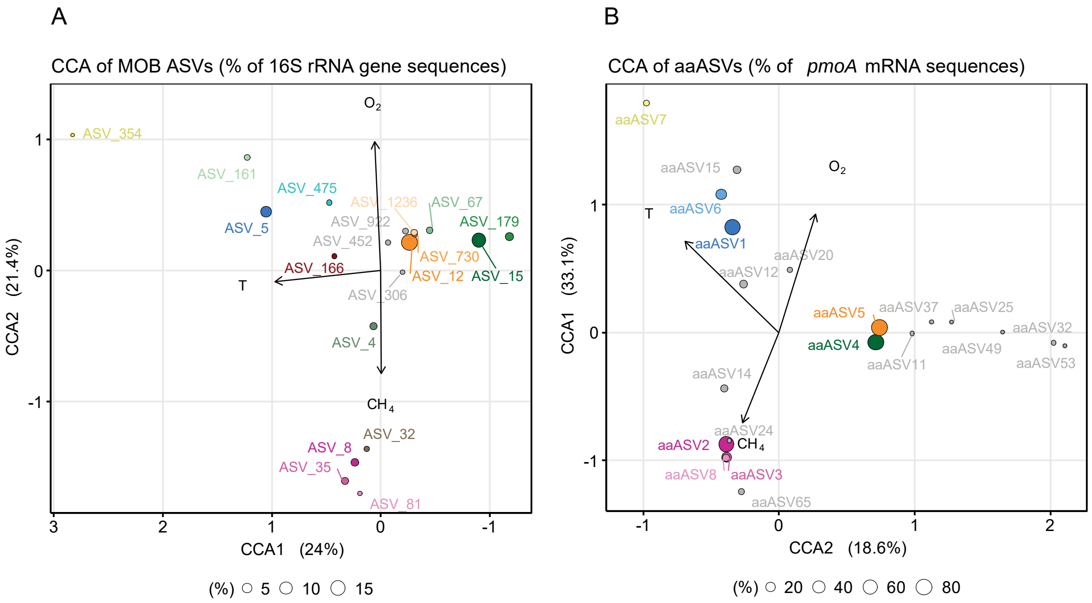


**Supplementary Figure 3** Canonical correspondence analysis (CCA) calculated based on a Chi-square dissimilarity matrix using the proportion of 16S rRNA gene sequences identified as MOB and physico-chemical parameters (temperature (T), oxygen ($O_2$) and methane ($CH_4$). The CCA was calculated for **A)** 16S rRNA gene MOB ASVs and **B)** *pmoA* mRNA aaASVs of all samples during lake overturn. Percentage of explained variance is given next to each axis. Colors are ASV/aaASV specific and correspond to the colours used in Figures 2 and 3 of the main text. The dot size visualizes the maximum percentage of the ASV/aaASV during lake overturn.

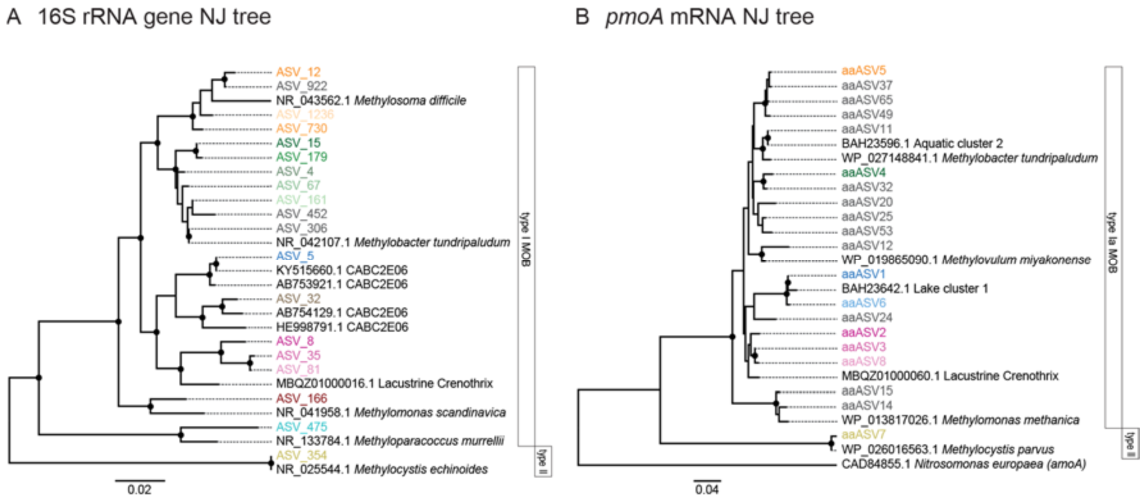


**Supplementary Figure 4** Phylogenetic placement of MOB 16S rRNA genes and *pmoA* amino acid sequences found in this study with reference sequences from cultivated and uncultivated MOB. Neighbor joining (NJ) trees were inferred with MEGA7 based on **A)** partial 16S rRNA gene sequences (423bp) and Jukes-Cantor evolutionary distance, and **B)** partial *pmoA* amino acid sequences (155 positions) and Poisson correction method. Black dots indicate nodes with a bootstrap value above 0.7 (10 000 bootstrap replicates). Colors are ASV/aaASV specific and correspond to the colours used in Figures 2 and 3 of the main text. Reference sequences are given with accession numbers to the left of the species name, or cluster name in case of uncultivated MOB. Scale bars represent changes per nucleotide or amino acid position.

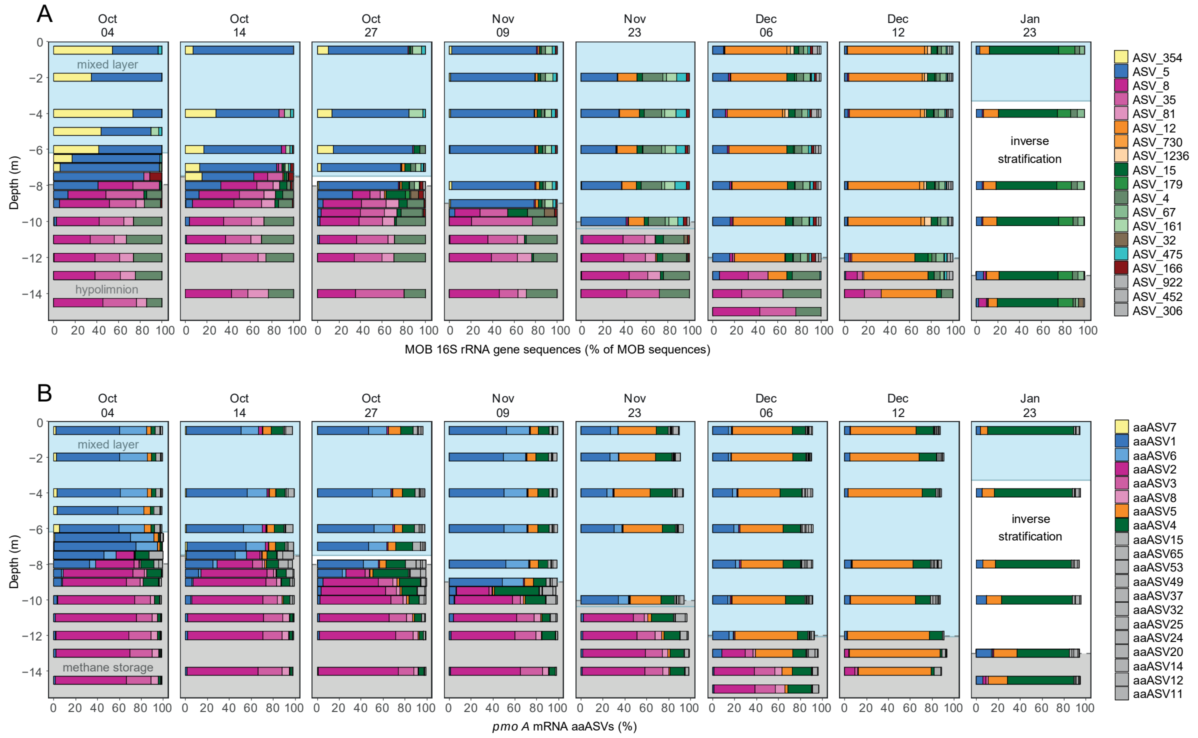


**Supplementary Figure 5** Depth distribution and dynamics within the MOB assemblage during lake overturn. The mixed layer depth increased with time (blue background), while the methane-rich bottom water (grey background) got gradually incorporated into the mixed layer. **A)** proportion of MOB ASVs among all identified MOB based on 16S rRNA gene amplicon sequencing **B)** Proportion of *pmoA* mRNA aaASVs based on amplicon sequencing. Colors are ASV/aaASV specific and correspond to the colours used in Figures 2 and 3 of the main text. ASVs and aaASVs which, based on multiple lines of evidence (see methods), are thought to originate from the same organism are encoded with the same color.

### Supplementary Tables

**Supplementary Table 1** Spearman rank correlation table of the MOB parameters and potential methane oxidation rate showing the strength of correlation. p-values were <0.001 after adjusting for multiple comparisons in R using the Benjamini & Hochberg (BH) method (Benjamini, Y., and Hochberg, Y. (1995). Controlling the false discovery rate: a practical and powerful approach to multiple testing. *Journal of the Royal Statistical Society Series B*, 57, 289–300).

| Spearman's rho | *pmoA* mRNA (copies $ml^{-1}$) | *pmoA* DNA (copies $ml^{-1}$) | MOB 16S rRNA gene (%) | MOB 16S rRNA (%) | MOB cells ($ml^{-1}$) |
|---|---|---|---|---|---|
| $CH_4$ oxidation rate (µM $d^{-1}$) | 0.82 (n=83) | 0.64 (n=83) | 0.88 (n=83) | 0.84 (n=83) | 0.75 (n=69) |
| *pmoA* mRNA (copies $ml^{-1}$) | | 0.83 (n=83) | 0.90 (n=83) | 0.81 (n=84) | 0.83 (n=68) |
| *pmoA* DNA (copies $ml^{-1}$) | | | 0.75 (n=83) | 0.62 (n=83) | 0.79 (n=68) |
| MOB 16S rRNA gene (%) | | | | 0.91 (n=83) | 0.82 (n=68) |
| MOB 16S rRNA (%) | | | | | 0.69 (n=68) |

**Supplementary Table *2*** Summary of measured mixed layer values used to produce Figure 3B and 3C. Abbreviations: Temp=Temperature, FCM = cells based on flow cytometry, pot. = potential, NA = missing values. Measurements smaller than the limit of quantification (LOQ) are indicated with "<" and the value of LOQ. In case of methane the lowest calibration concentration is shown. LOQ of nitrate was derived from the FIA instrument baseline.

| Date DD.MM.YYYY | Layer | Depth (m) | Temp (°C) | Oxygen (µmol $l^{-1}$) | Methane (µmol $l^{-1}$) | Nitrate (µmol $l^{-1}$) | FCM (cells $ml^{-1}$) | Pot. methane oxidation rate (µmol $l^{-1}$ $d^{-1}$) |
|---|---|---|---|---|---|---|---|---|
| 04.10.2016 | mixed | 0.50 | 18.4 | 413 | 0.3 | < 4 | 6.3E+06 | NA |
| 04.10.2016 | mixed | 2.00 | 18.5 | 412 | 0.3 | < 4 | 5.8E+06 | 0.2 |
| 04.10.2016 | mixed | 4.00 | 18.5 | 412 | 0.3 | < 4 | 5.4E+06 | 0.1 |
| 04.10.2016 | mixed | 5.00 | 18.4 | 412 | 0.2 | < 4 | 5.7E+06 | 0.2 |
| 04.10.2016 | mixed | 6.00 | 18.4 | 412 | 0.3 | < 4 | 5.9E+06 | 0.2 |
| | **MEDIAN:** | **n=5** | **18.4** | **412** | **0.3** | **< 4** | **5.8E+06** | **0.2** |
| 14.10.2016 | mixed | 0.50 | 14.4 | 279 | 0.6 | < 4 | 6.2E+06 | 0.5 |
| 14.10.2016 | mixed | 4.00 | 14.4 | 276 | 0.6 | < 4 | 6.2E+06 | 0.6 |
| 14.10.2016 | mixed | 6.00 | 14.4 | 277 | <0.05 | < 4 | 6.1E+06 | 0.4 |
| 14.10.2016 | mixed | 7.00 | 14.4 | 273 | 0.7 | < 4 | 6.0E+06 | 0.5 |
| | **MEDIAN:** | **n=4** | **14.4** | **277** | **0.6** | **< 4** | **6.1E+06** | **0.5** |
| 27.10.2016 | mixed | 0.50 | 13.1 | 290 | 0.1 | < 4 | 5.2E+06 | 0.3 |
| 27.10.2016 | mixed | 4.00 | 13.0 | 287 | 0.1 | < 4 | 5.8E+06 | 0.7 |
| 27.10.2016 | mixed | 6.00 | 13.0 | 285 | 0.2 | < 4 | 5.2E+06 | 0.5 |
| 27.10.2016 | mixed | 7.00 | 13.0 | 280 | 0.2 | < 4 | 5.3E+06 | 1.0 |
| | **MEDIAN:** | **n=4** | **13.0** | **286** | **0.1** | **< 4** | **5.3E+06** | **0.6** |
| 09.11.2016 | mixed | 0.50 | 10.7 | 272 | 1.4 | 4.6 | 5.6E+06 | 2.5 |
| 09.11.2016 | mixed | 2.00 | 10.7 | 271 | 1.3 | 5.2 | 5.4E+06 | 2.5 |
| 09.11.2016 | mixed | 4.00 | 10.7 | 270 | 1.0 | 5.4 | 5.5E+06 | 2.4 |
| 09.11.2016 | mixed | 6.00 | 10.7 | 269 | 1.0 | 5.6 | 5.4E+06 | 3.0 |
| 09.11.2016 | mixed | 8.00 | 10.7 | 270 | 1.1 | 6.1 | 5.2E+06 | 2.6 |
| | **MEDIAN:** | **n=5** | **10.7** | **270** | **1.1** | **5.4** | **5.4E+06** | **2.5** |
| 23.11.2016 | mixed | 0.50 | 8.9 | 248 | 0.1 | 8.0 | NA | 2.2 |
| 23.11.2016 | mixed | 2.00 | 8.9 | 245 | 0.1 | 8.5 | NA | 2.5 |
| 23.11.2016 | mixed | 4.00 | 8.9 | 244 | 0.1 | 8.4 | NA | 2.8 |
| 23.11.2016 | mixed | 6.00 | 8.9 | 245 | <0.05 | 8.3 | NA | 2.4 |
| 23.11.2016 | mixed | 8.00 | 8.9 | 243 | <0.05 | 8.4 | NA | 2.7 |
| | **MEDIAN:** | **n=5** | **8.9** | **245** | **0.1** | **8.4** | **NA** | **2.5** |
| 06.12.2016 | mixed | 0.50 | 7.0 | 181 | 0.4 | 8.2 | 2.5E+06 | 5.3 |
| 06.12.2016 | mixed | 2.00 | 7.0 | 179 | 0.7 | 8.5 | 2.5E+06 | 5.3 |
| 06.12.2016 | mixed | 4.00 | 7.0 | 179 | 0.7 | 8.7 | 2.5E+06 | 5.6 |
| 06.12.2016 | mixed | 6.00 | 7.0 | 180 | 0.5 | 9.1 | 2.5E+06 | 5.4 |
| 06.12.2016 | mixed | 8.00 | 7.0 | 178 | 0.9 | 8.7 | 2.5E+06 | 5.8 |
| 06.12.2016 | mixed | 10.00 | 7.0 | 177 | 0.5 | 8.7 | 2.5E+06 | 5.2 |
| | **MEDIAN:** | **n=6** | **7.0** | **179** | **0.6** | **8.7** | **2.5E+06** | **5.3** |
| 12.12.2016 | mixed | 0.50 | 6.1 | 175 | <0.05 | 8.8 | 2.1E+06 | 7.2 |
| 12.12.2016 | mixed | 2.00 | 6.1 | 175 | <0.05 | 8.8 | 2.1E+06 | 6.4 |
| 12.12.2016 | mixed | 4.00 | 6.1 | 173 | <0.05 | 9.3 | 1.9E+06 | 6.2 |
| 12.12.2016 | mixed | 8.00 | 6.1 | 175 | 0.3 | 9.1 | 2.1E+06 | 7.8 |
| 12.12.2016 | mixed | 10.00 | 6.0 | 172 | <0.05 | 9.0 | 1.9E+06 | 6.3 |
| | **MEDIAN:** | **n=5** | **6.1** | **175** | **<0.05** | **9.0** | **2.1E+06** | **6.4** |
| **23.01.2017** | **mixed** | **0.50** | **1.4** | **329** | **0.1** | **10.6** | **5.0E+06** | **2.0** |
| | **MEDIAN:** | **n=1** | | | | | | |
